# Theory and practice of using cell strainers to sort *Caenorhabditis elegans* by size

**DOI:** 10.1101/2023.01.07.523116

**Authors:** Vincent J. Lanier, Amanda M. White, Serge Faumont, Shawn R. Lockery

## Abstract

The nematode *Caenorhabditis elegans* is a model organism widely used in basic, translational, and industrial research. *C. elegans* development is characterized by five morphologically distinct stages, including four larval stages and the adult stage. Stages differ in a variety of aspects including size, gene expression, physiology, and behavior. Enrichment for a particular developmental stage is often the first step in experimental design. When many hundreds of worms are required, the standard methods of enrichment are to grow a synchronized population of hatchlings for a fixed time, or to sort a mixed population of worms according to size. Current size-sorting methods have higher throughput than synchronization and avoid its use of harsh chemicals. However, these size-sorting methods currently require expensive instrumentation or custom microfluidic devices, both of which are unavailable to the majority *C. elegans* laboratories. Accordingly, there is a need for inexpensive, accessible sorting strategies. We investigated the use of low-cost, commercially available cell strainers to filter *C. elegans* by size. We found that the probability of recovery after filtration as a function of body size for cell strainers of three different mesh sizes is well described by logistic functions. Application of these functions to predict filtration outcomes revealed non-ideal properties of filtration of worms by cell strainers that nevertheless enhanced filtration outcomes. Further, we found that serial filtration using a pair of strainers that have different mesh sizes can be used to enrich for particular larval stages with a purity close to that of synchronization, the most widely used enrichment method. Throughput of the cell strainer method, up to 14,000 worms per minute, greatly exceeds that of other enrichment methods. We conclude that size sorting by cell strainers is a useful addition to the array of existing methods for enrichment of particular developmental stages in *C. elegans*.

## Introduction

The nematode *Caenorhabditis elegans* is a model organism widely used in basic, translational, and industrial research. It can be grown quickly (three-day generation time) and maintained in large numbers (tens of thousands) at negligible cost relative to zebrafish and rodents. It has the most comprehensively annotated genome of all model organisms. The entire set of cell divisions from oocyte to the 995 somatic cells of the mature worm has been mapped. Of the somatic cells, 302 are neurons, and their anatomical connectivity has been described completely, a first in any organism. The *C. elegans* genome exhibits remarkably strong homologies to mammals including humans. It is estimated that 60–80% of *C. elegans* genes have a homolog in humans [1]. Phenotypes of numerous human diseases have been replicated in *C. elegans* by manipulating these genes [2]. The worm also exhibits a range of complex, experience-dependent behaviors that provide models for similar behaviors in mammals with much larger brains [3,4]. These strengths are coupled with unsurpassed genetic tractability, including a substantial molecular biological toolkit that accelerates research at multiple levels of organization, from genes to cells and behavior.

*C. elegans* development is characterized by five morphologically distinct stages, including four larval stages (L1–L4) and the adult stage, each separated by an observable molt. Stages differ in a variety of aspects including size, gene expression, physiology, and behavior. Thus, enrichment of a particular developmental stage is often the first step in experimental design. When several hundred worms are required, the method of enrichment used in most laboratories is to culture a developmentally synchronized population of hatchlings for a fixed time. This is accomplished by dissolving gravid adults in a caustic bleach solution to collect eggs, which remain viable (henceforth *bleach synchronization*). Hatchlings are synchronized by maintaining them overnight in the absence of food, causing them to enter developmental arrest at the L1 stage. It is possible that bleaching worms can have multigenerational detrimental effects on the physiology and behavior of progeny [5]. Moreover, bleaching cannot be used to enrich for particular larval stages after an experimental treatment such as application of a drug or changes in diet. The main alternative is to sort worms by size. This approach is advantageous because it does not require bleaching, and it generally has higher throughput. The highest throughput and purity is achieved by the COPAS Biosort (Union Biometrica, Holliston, MA, USA), a flow cytometer that accommodates *C. elegans* [6]. Though highly effective, the high cost of the Biosort limits its accessibility to most laboratories. Size sorting has the limitation that mutations and drug treatments can change worm dimension such that sorting protocols may require compensatory adjustments.

In response to the drawbacks of synchronization and the Biosort, a variety of microfluidic devices (chips) have been developed that sort worms by size, behavior, or physical properties. These include planar microsieves [7–12], chips that accentuate stage-specific behavioral responses to an applied electric field [13–16], inertial [17] or acoustic focusing [18], and impedance measurements [19]. However, because sorting chips are not commercially available, they must be made by the user, a process that requires expensive facilities and expertise in microfabrication. Indeed, cited-reference searches reveal that adoption by *C. elegans* laboratories of sorting chips published by bioengineering researchers is absent or extremely limited.

Accordingly, there is a need for simple, inexpensive sorting strategies that avoid the use of bleach, provide large yields at high-throughput, and utilize readily available materials. We investigated the use of mesh filters, such as commercially available cell strainers, to sort *C. elegans* by size. A cell strainer is a small plastic cup the bottom of which is a nylon mesh that has a well-defined opening size. Cell strainers are available in a wide range of mesh sizes, from 1 to 1000 μm.

Cell strainers are capable of separating very young worms (L1s, L2s) from mixed-stage populations [20,21]. Here we demonstrated the use of mesh filters to recover later developmental stages. We found that recovery probability for cell strainers of three different mesh sizes is well described by logistic functions. Application of these functions to predict filtration outcomes revealed non-ideal properties of filtration by cell strainers that nevertheless enhanced filtration outcomes. Additionally, we found that serial filtration using a pair of strainers of different mesh sizes can be used to enrich for particular larval stages with a purity approaching that of synchronization, the most widely used enrichment method. Throughput of the cell-strainer method was 9,000 – 14,000 worms per minute, greatly exceeding that of all other enrichment methods. We conclude that size sorting by cell strainers is a useful addition to the array of existing methods for enrichment of particular developmental stages of *C. elegans*.

## Materials and Methods

### Nematodes

The reference strain of *C. elegans* (N2) was obtained from the *Caenorhabditis* Genetics Center at the University of Minnesota (St. Paul). Worms were grown at 20 C on 50 mm petri plates filled with NGM agarose and seeded with the OP50 strain of *E. coli* [22]. Plates of mixed-stage worms were obtained by transferring small cubes of worm-laden agarose (5 mm × 5 mm × 5 mm) to the plates and allowing cultures to grow until most of the *E. coli* were consumed (3 days).

### Solutions

M9 buffer was used to wash worms off culture plates and to suspend worms during filtration. This buffer was prepared by combining 3 g KH2 PO4, 6 g Na2 HPO4, 5 g NaCl and 1 mL of 1 M MgSO4, and adding H2O to 1 L.

### Synchronization

Worms were washed off a culture plate carrying a mixed-stage population of worms, including an ample number of gravid hermaphrodites. Worms were recovered into 15 mL centrifuge tubes, then pelleted by centrifugation at 2000 rpm for one minute. The supernatant was discarded, and then 1 mL of household bleach (8.25% solution of sodium hypochlorite) and 200 μL 5M KOH were added. While the tube was agitated, the bleaching progress was observed under a microscope. The reaction was stopped just before all traces of worm bodies disappeared, leaving only eggs behind (about 4 minutes). Tubes were topped off with M9 and centrifuged at 2000 RPM for one minute, followed by two additional M9 washes. Eggs were then transferred to foodless culture plates and allowed to hatch overnight at 20 C. The following day, hatchlings (L1) were washed off the unseeded plates with M9, recovered in 15 mL falcon tubes, and transferred to seeded *E. coli* plates.

### Filtration apparatus

Cell strainers and accessories were purchased from PluriSelect USA, El Cajon, CA, USA. The filtration system comprised a nested stack of two cell strainers wherein the upper strainer had a larger mesh size than the lower strainer (Fig 1). A 25 mL loading funnel was inserted into the upper strainer. The lower strainer was inserted into an adapter which was inserted into the mouth of a stack of two 50 mL centrifuge tubes. The conical bottom of the upper tube was cut off to enable this tube to be inserted into the lower tube. The joint between tubes was sealed with tape or heat-shrink tubing.

**Fig 1.**
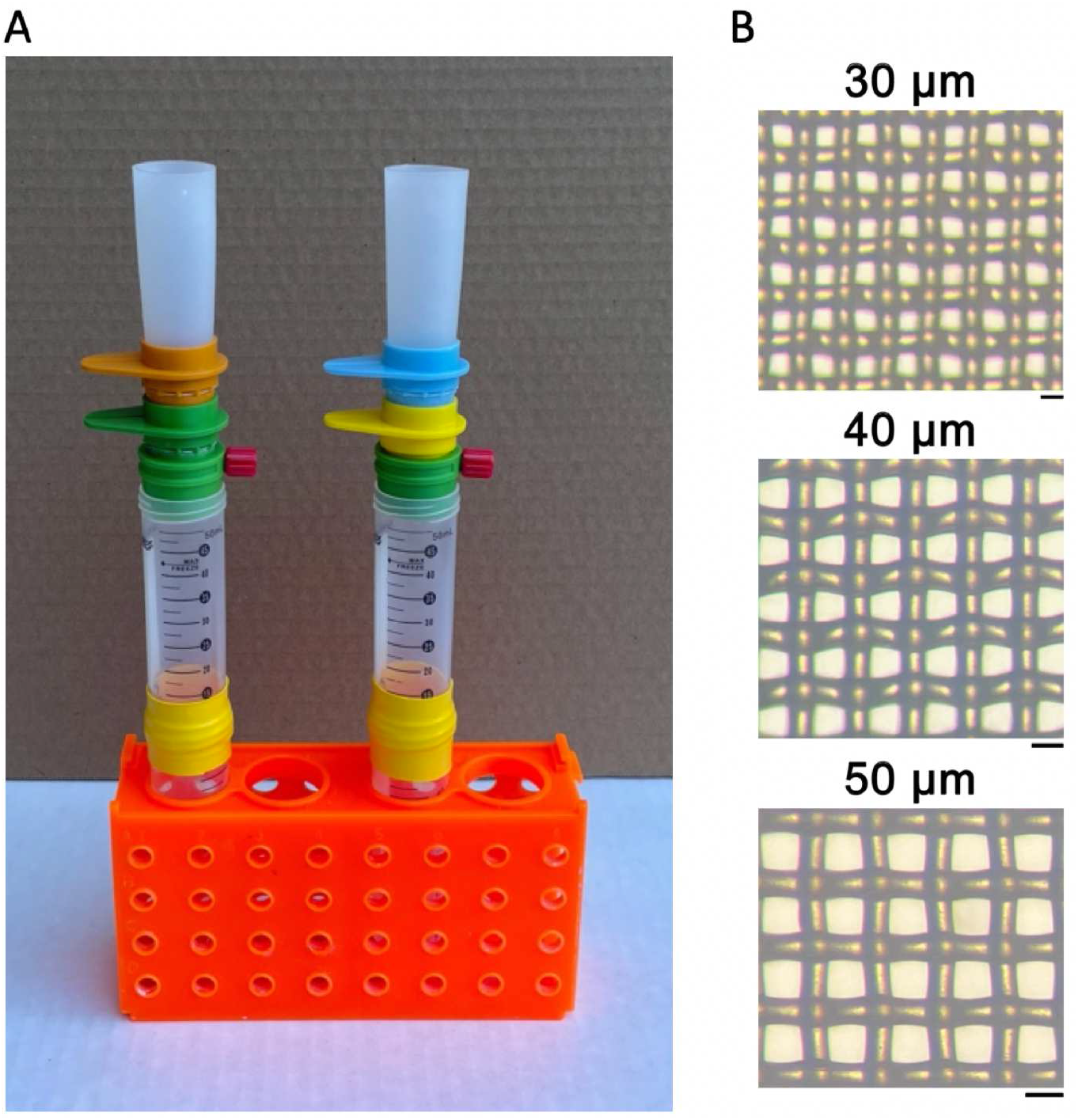
Filtration apparatus. A. Left, strainer stack for pre-filtration. An 85 μm strainer (orange) is stacked on a 20 μm strainer (green). A funnel is inserted into the 85 μm strainer and the 20 μm strainer is inserted into an adapter whose vent port is closed by a red cap. Right, strainer stack for 1-step passage, 40 μm filtration. A 40 μm strainer (blue) is stacked on a 5 μm strainer (yellow). B. Micrographs of the mesh in 30, 40, and 50 μm strainers. Scale-bars equal mesh size.

### Pre-filtration

The purpose of this step was to remove debris and eliminate most of the of L1 and L2 worms, which were not part of this study. An 85 μm cell strainer was stacked on a 20 μm strainer (85/20). A vent port on the side of the adapter could be closed by a cap to prevent liquid from flowing out of the funnel or opened to initiate the filtration process. Approximately 2 mL of M9 buffer was added to each plate to suspend worms in fluid. Each plate was emptied into the funnel, then rinsed with a stream of M9 from a squirt bottle to transfer any remaining worms. The funnel was then topped up with M9 after which the vent port was opened and fluid was allowed to drain through the filters. Any fluid remaining in the top strainer was removed by applying gentle suction via a 10 mL syringe attached to the vent port. The port was again closed, and the funnel was re-filled with M9 and drained again. This process was repeated for a total of three *washes* of the funnel; each set of 3 washes is referred to as a single filtration *step*. To recover worms from the 20 μm strainer, the stack was disassembled and the 20 μm strainer was inverted over a glass funnel (40 mm diameter) inserted into a 10 mL centrifuge tube. Worms were recovered from the strainer by back flushing with a stream of M9 until no worms remained on the strainer, determined by examination under a stereomicroscope. As described in Results, pre-filtered worms were divided into populations *P*_1_ and *P*_2_ for subsequent filtration. This was done in one of two ways: (i) by resuspending pre-filtered worms by agitation of the 10 mL centrifuge tube, then placing half of the fluid in each of two 10 mL centrifuge tubes, or (ii) by pelleting the pre-filtered worms by settlement (5 min at −20 C), sampling *P*_1_ directly from the pellet, and resuspending the worms remaining in the pellet to form *P*_2_. The two methods are functionally the same.

### Size sorting

The size-sorting filtration protocols were similar to pre-filtration, including assembly of the filter system, transfer of worms from plates into the funnel, the number of washes in each filtration step, and the method for recovering worms from the strainer. The population of worms recovered after filtration was designated *R*. There were two main differences between pre-filtration and size-sorting protocols. (i) The mesh sizes of strainers in the filter stack were altered as described in Table 1 and Fig 2. (ii) A 10 mL syringe was attached to the vent port to regulate fluid flow from filters to the collection tube. Regulation was necessary because when filters in the range of 30 – 50 μm were used, flow rates varied according to the number of worms in the funnel and the mesh size. Using the syringe to form a modest vacuum, we adjusted flow rate so that fluid height in the funnel dropped by approximately 2–3 mm/sec. Some experiments involved a single filtration step, whereas others involved two or three steps (Fig 2). Each filtration experiment was replicated at least 7 times; each replicate is called a *run*. After use, strainers were cleaned by sonication in distilled water, or by immersion in soapy water, rinsed, and dried with a stream of air. Strainers cleaned in this way could be used indefinitely.

**Table 1.** Filtration protocols, number of filtration runs, and figures in which the associated data appear.

| Filtration method | Number of steps | Number of plates/run | Number of runs | Figure |
| --- | --- | --- | --- | --- |
| Pre-filtration | 1 | – | – | – |
| 1-step retention, 30 $\mu\text{m}$ | 1 | 4–7 | 8 | 3A |
| 1-step retention, 40 $\mu\text{m}$ | 1 | 4–7 | 8 | 3B |
| 1-step passage, 40 $\mu\text{m}$ | 1 | 4–7 | 11 | 3C |
| 1-step passage, 50 $\mu\text{m}$ | 1 | 4–7 | 9 | 3D |
| 3-step retention, 40 $\mu\text{m}$ | 3 | 9–10 | 8 | 7B |
| L4-YA Method 1 | 2 | 6–8 | 7 | 8A |
| L4-YA Method 2 | 1 | 10 | 7 | 8B |
| L4-YA Method 3 | 2 | 10 | 7 | 8C |

**Fig 2.**
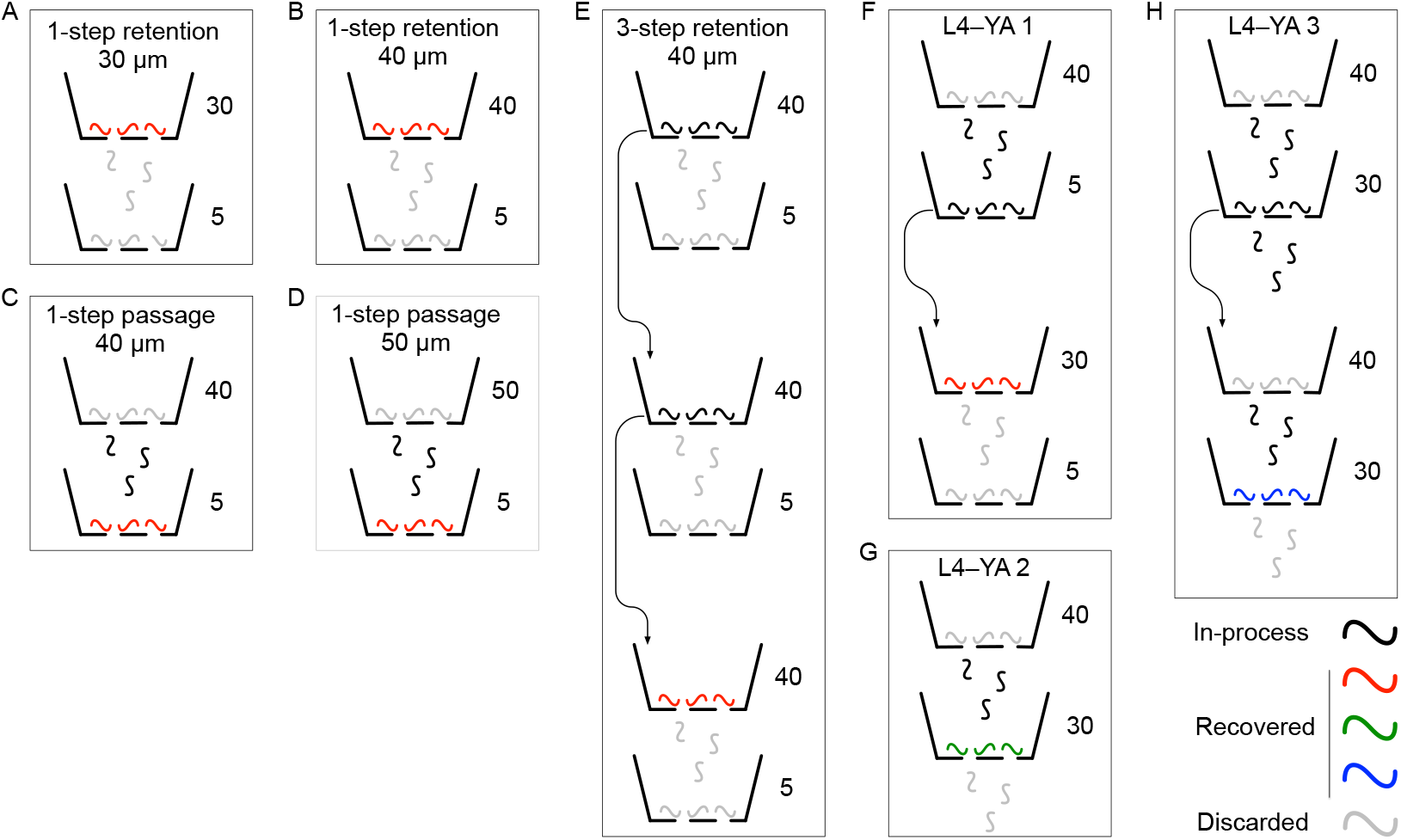
Flow charts of the eight size-sorting filtration protocols tested in this study. Numbers next to strainer icons indicate mesh size (μm). Worm symbols are color coded according to phase in the filtration process as shown in the key. *Arrows* indicate successive filtration steps, as defined in Materials and Methods. Although not strictly necessary, the 5 μm filters in 1-step retention, 30 μm and 1-step retention, 40 μm were included for consistency with the 1-step passage, 40 μm and 1-step passage, 50 μm protocols.

### Measurement of worm length

Worm lengths in *P*_1_ and *R* were measured by imaging worms on 50 mm, bacteria-free NGM plates. After pelleting by settlement (5 min at −20 C) worms were transferred to the plates in 2– 5 μL aliquots until the desired number of worms was obtained (usually, 3–6 aliquots per plate). Prior to measurement, worms were allowed to disperse for 5–10 min., a process sometimes accelerated by puffs of air from a rubber bulb. Typically, all worms on 4–6 plates were imaged per experiment. Worms were immobilized by flowing C02 through a transparent chamber inverted over the plate. Worms were imaged at 5.64 μm/pixel using a macro lens (AF Micro-Nikkor 60 mm f/2.8D, Nikon, Japan). A series of still images tiling the entire surface of the plate was captured using a microscope camera (HDMI 1080P HD212, AmScope, Irvine, CA, USA). Images were combined into a single image stack which was submitted to WormLab software (MBF Bioscience, Williston, VT, USA) for automated length measurements. Length measurements excluded the worm’s tail which was not resolved in the images.

### Developmental stages and dimensions

The plot in S1 Fig shows the relationship between width and length used in this study. Widths of each developmental stage were taken from Table 1 of Atakan et al. [7]. Lengths of each stage were the center of the length ranges that define each stage according to data in WormAtlas [23]. Plotting width (*w*, μm) against length (*l*, μm) yielded a linear relationship between L1 and YA, with *w* = 0.055*l* – 3.53.

### Estimation of throughput

Throughput was defined as the number of worms filtered per minute of processing time. To estimate the number of worms that were loaded into the filtration apparatus per culture plate, a suspension of worms was obtained by washing each of 6 typical, mixed-stage culture plates with 2 mL M9 buffer. Worms were pelleted by settlement (5 min at −20 C) after which the volume of the suspension was adjusted to 5 mL. The pellet was resuspended by agitation. Resuspended worms were transferred to 11 foodless NGM plates (20 μL per plate). Worms were allowed to disperse for approximately 10 min, then counted in images of the plates taken on a flat-bed scanner (Epson V850 Pro, Los Alamitos, CA, USA). This procedure yielded an estimated worm density of 5.6 worms/μL in the 5 mL suspension, which is equivalent to 4.7 × 10^3^ worms/plate. Throughput estimates were based on a typical value of 10 plates per filtration run. Throughput was computed as *N/T_i_* where *N* is the estimated number of worms on 10 plates (4.7 × 10^4^), and *T_i_* is processing time of run *i* which began at the moment the contents of the first culture plate were loaded into the pre-filtration filter stack and ended when worms were recovered from the filtration stack. Throughput values in Table 2 are within-method mean processing time (*n* = 3 runs). Processing time *T* for bleach synchronization included culture time (48 hr.).

**Table 2.** Performance metrics of filtration methods for enrichment of particular larval stages.

| Parameter | Method 1 | Method 2 | Method 3 | Synch |
| --- | --- | --- | --- | --- |
| Total number of worms measured | 3545 | 6333 | 5793 | 2175 |
| Grand mean of run means (length, $\mu\text{m}$) | $872 \pm 59$ | $778 \pm 53$ | $773 \pm 27$ | $812 \pm 35$ |
| Mode<sup>a</sup> ($\mu\text{m}$) | 825 | 725 | 775 | 825 |
| Gaussian means ± std. dev. in Fig 8 ($\mu\text{m}$) | $867 \pm 159$ | $754 \pm 165$ | $768 \pm 141$ | $826 \pm 101$ |
| Mean processing time (min) | $6.4 \pm 0.1$ | $3.3 \pm 0.2$ | $5.2 \pm 0.5$ | 2880<sup>b</sup> |
| Mean throughput (worms/min) | - | $14,100^c \pm 690$ | $9,100^c \pm 810$ | 1.63<sup>d</sup> |
| Mean yield | - | $1,130^e \pm 160$ | $755^e \pm 260$ | $242^f \pm 77$ |
| Mean purity<sup>g</sup> | $0.35 \pm 0.14$ | $0.45 \pm 0.079$ | $0.54 \pm 0.076$ | $0.65 \pm 0.085$ |

Metrics for synchronized worms are provided for comparison. Means are shown ± std. dev. <sup>a</sup>Most frequent worm length. <sup>b</sup>L1 larvae were grown for 48 hr. <sup>c</sup>Based on ten 5 cm plates of worms washed onto the funnel. <sup>d</sup>Based on the estimated number of worms in one mixed stage culture plate (Materials and Methods) <sup>e</sup>Lower-bound yield. <sup>f</sup>Yield from bleaching one mixed-stage culture plate. <sup>g</sup>The purity of unfiltered worms was $0.191 \pm 0.08$ (std. dev.).

### Measurement of sorting purity

Purity was defined as the proportion of worms in filtered distributions whose lengths were within the target range of developmental stages, L4–YA. Purity on each run was computed as the integral within the target zone of probability densities *H*_u_(*l*) and *H*_f_(*l*). Purity for synchronized populations was computed in a similar manner, with one exception. The peak of the mean probability density function for synchronized worms fell within the target zone but was not centered on it. Therefore, before computing purity values for each synchronization each run, we centered its histogram by shifting it 39 μm to the left. This approach eliminated the spurious reduction in purity that would have resulted from the slight mismatch between the synchronized distribution and the target range.

### Statistical analyses

Two-tailed *t*-tests were used to compare sorting purities between filtered and unfiltered populations, and between populations filtered by different methods. Degrees of freedom *v* in *t*-tests were computed, without the assumption of equal variances, as

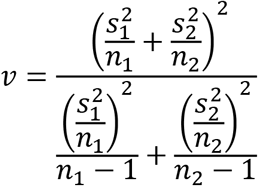

where *s_i_* is the standard deviation of purity for method *i* and *n_i_* is the number of runs for that method. Statistical tests were implemented in Igor Pro (Wavemetrics, Portland, OR, USA).

## Results

Previous studies used filtration through commercially available cell strainers as a means of enriching for worms in the first larval stage (L1), or the first and second larval stages (L1, L2), from mixed-stage populations [20, 21]. We tested the ability of cell strainers to enrich for later stages, including L4s and young adults (YA). To ensure that the filtration methods developed here could be integrated easily into common workflows, we adopted widely used methods to create mixed-stage cultures on standard agarose plates (Materials and Methods). At the start of each filtration experiment, worms were washed off culture plates and pre-filtered to remove debris and L1–L2 stage worms. This was done using an 85 μm cell strainer stacked on a 20 μm cell strainer (85/20). Worms recovered from the 20 μm filter were then subjected to one of eight different filtration protocols selected according to the purpose of the experiment (Fig 2).

### Mathematical models of filtration

We started with the simple case of 1-step filtration using a single cell strainer (Fig 2A-D, Fig 3A-D). For each run, pre-filtered worms were divided into separate populations, *P*_1_ and *P*_2_. The lengths of worms in a subsample of *P*_1_ were measured to construct a histogram estimating the length distribution of the worms to be loaded into the funnel of the filtration setup. We refer to the histogram of *P*_1_ as the *unfiltered* distribution, *H*_u_(*l*). Unfiltered distributions varied between experiments, but there was a consistent bias toward earlier developmental stages (Fig 3, black traces), which reflects the particular worm culturing protocol we used (Methods and Materials). Population *P*_2_ was loaded into the funnel, from which we recovered the population *R*, containing size-sorted worms. The lengths of worms in a subsample of *R* were measured to construct a histogram estimating the length distribution of *filtered* worms, *H*_f_(*l*). These histograms, converted to units of probability density and averaged across runs, and named 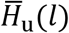 and 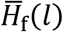, are shown in Fig 3. An empirically derived length-stage relationship was used to infer which developmental stages were obtained in 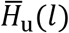 and 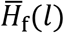 (Materials and Methods and S1 Fig).

**Fig 3.**
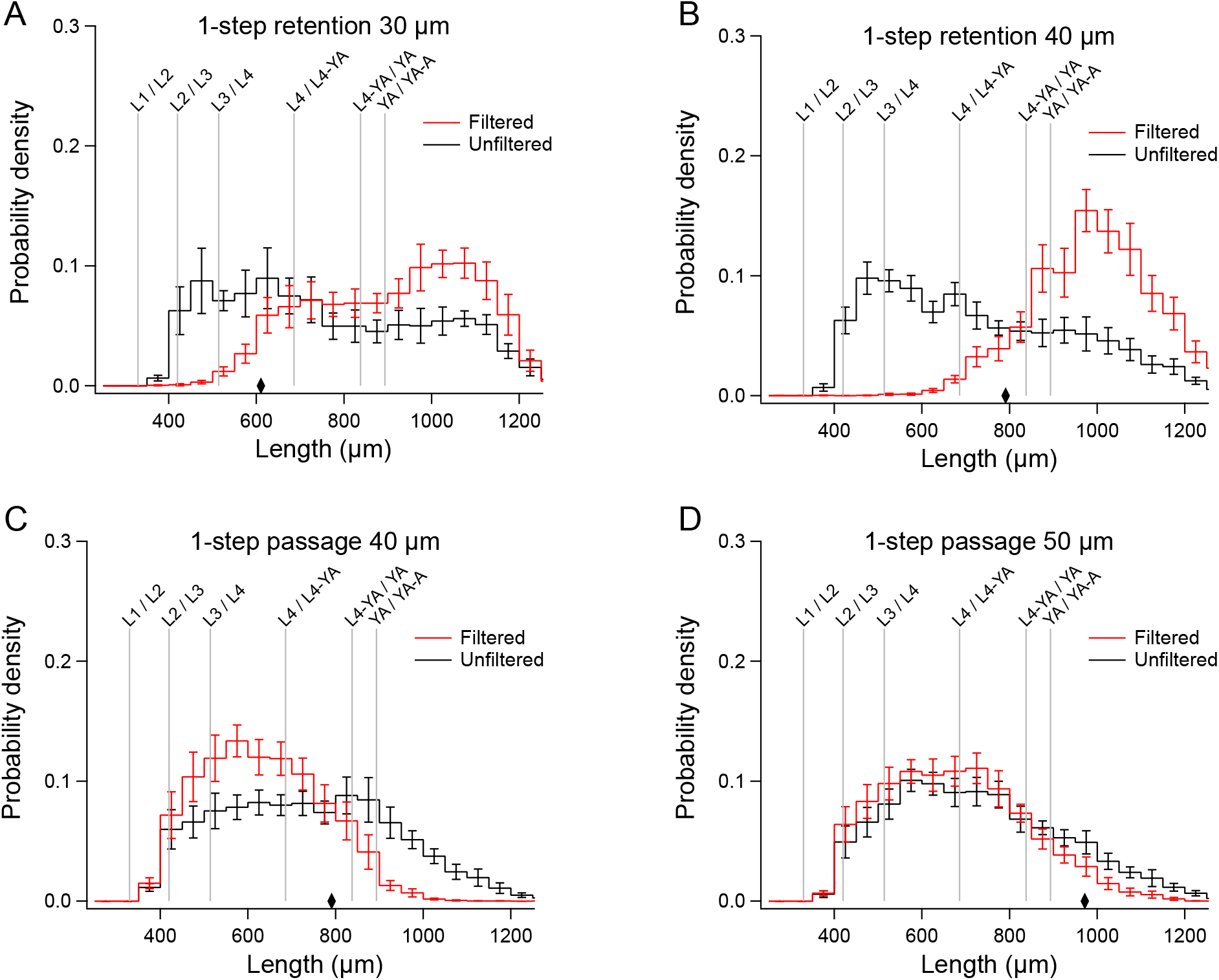
Mean distributions of filtered and unfiltered worms in 1-step filtration experiments. A-D. Mean filtered distributions, 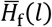 (*red traces*) and mean unfiltered distributions, 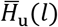 (*black traces*). Diamonds indicate the length of worms having a diameter equal to the mesh size. Error bars, SEM.

Like any sieve, a cell strainer can be used in two *modes*, retention and passage. The distribution of worms recovered from the top surface of the strainer (*retained fraction*) was shifted toward larger worms relative to the unfiltered distribution, whereas the length distribution of worms recovered from the fluid that passed through the filter (*passed fraction*) was shifted to toward smaller worms (Fig 3). This pattern indicates that cell strainers used in our experiments (30 μm, 40 μm, and 50 μm) can be used to obtain populations enriched for larger or smaller worms in the length range 400 – 1200 μm (L3 and older).

The diamonds in Fig 3 indicate the length of worms having a width equal to the mesh size (S1 Fig). In retention graphs, non-zero points to the left of the diamonds indicate worms that were retained even though they were narrower than the mesh size. We hypothesize that retention of narrower worms occurs when worms land on the strainer mesh roughly parallel it. In passage graphs, non-zero points to the right of the diamonds signify worms that were wider than the mesh size, but nevertheless passed through the filter. The passage of wider worms might be explained by deformation by fluid pressure as they pass through the filter.

Any sieve has a characteristic *filter function*, defined as the probability *p* that a particle will be recovered as a function of its size, *x*. This probability is computed as

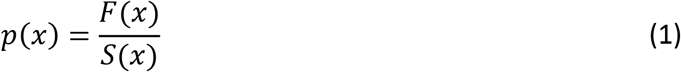

where *F*(*x*) is the number of recovered particles in bin *x* and *S*(*x*) is the number of particles in bin *x* in the starting distribution. The filter function is useful because it is independent of the starting distribution of particle sizes, making it a universal statement of filter selectivity. Moreover, it can be used to predict filtration outcomes with respect to any starting distribution simply by bin-wise multiplication of the filter function and the starting distribution; such predictions can facilitate design of custom filtration protocols. In contrast, the filtered distribution is strongly dependent on the starting distribution: the greater the number of particles in a given bin in the starting distribution, the greater the number of recovered particles in the corresponding bin the filtered distribution (except where *p*(*x*) = 0).

Given the advantages of filter functions, we wished to recover these functions for the cell strainers used in this study. In this instance, equation (1) becomes

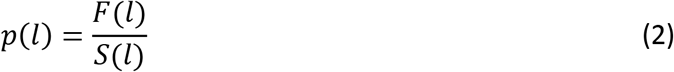

where *F*(*l*) is the raw length histogram of all worms recovered after filtration (population *R*) and *S*(*l*) is the raw length histogram all worms loaded into the funnel (population *P*_2_). In retention filtration, *p*(*l*) approaches unity in the limit of large *l*; in passage filtration, *p*(*l*) approaches unity in the limit of small *l*. To compute *p*(*l*) exactly, it would be necessary to measure the lengths of *all* worms in the *R* and *P*_2_ populations. However, because our filtration methods accommodate very large numbers of worms, it was not possible to measure the length of every worm. We relied instead on samples of these populations (Materials and Methods, Measurement of worm length). Nevertheless, given that our sampling method is unbiased, the forms of *H*_f_(*l*) and *H*_u_(*l*) should closely resemble the forms of *F*(*l*) and *S*(*l*), respectively. Moreover, their ratio should have the same form as *p*(*l*). However, because of sampling, the relative amplitudes of *H*_f_(*l*) and *H*_u_(*l*) will almost certainly differ from the relative amplitudes of *F*(*l*) and *S*(*l*), so the limiting values of *p*(*l*) will generally not be unity. Nevertheless, the expected limiting values of *p*(*l*) provided a constraint that we used to correct for sampling. In particular, we computed

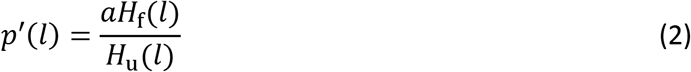

where *a* is a sampling correction factor whose value was chosen to ensure the expected limiting behavior of *p*(*l*). Specifically, we set *a* = 1/*X_n_*, the reciprocal of the mean of the *n* extremes values of *H*_f_(*l*)/*H*_u_(*l*) for each run. The value of *n* was selected to optimize the fitting procedure described immediately below; the sampling correction procedure is illustrated in S2 Fig.

The dependent variable in our filtration data is the probability of a favorable *binary, categorical* outcome (recovery of a worm from the filtration process, vs. failure of recovery). The independent variable is worm length. The the standard equation for relating the probability of binary outcomes to a continuous independent variable is called the *logistic function*. The standard statistical method for analyzing binary outcomes is *logistic regression*[24] which entails fitting the general form of the *logistic equation* to a specific data set. The logistic equation is an exponential sigmoid

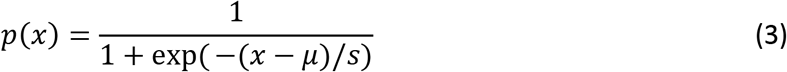

where *x* is the independent variable, *μ* is a location parameter (the point on the *x* axis where *p*(*x*) = 0.5), and *s* is a scale parameter. It ranges from zero at large negative values of (*x* – *μ*) to unity at large positive values of (*x* – *μ*). The logistic equation is rotationally symmetric about the point (μ, 0.5). In its rotated form, it ranges from unity to zero.

We found that after correction for sampling, mean retention data (Fig 4A,B, black symbols) were well fit by a logistic function having the form

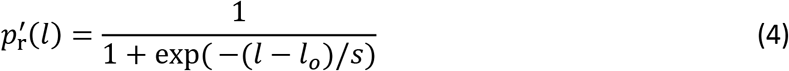

where *μ* = *l_o_*, the length at which the *p′*(*l*) = 0.5, (Fig 4A,B, red traces). Mean passage data were well fit by the rotated from of equation (4)

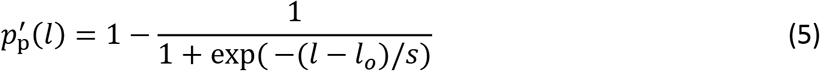

(Fig 4B,C). Parameters for each fitted function are given in S1 Table, including the value of *n* used to determine the sampling correction factor *a*.

**Fig 4.**
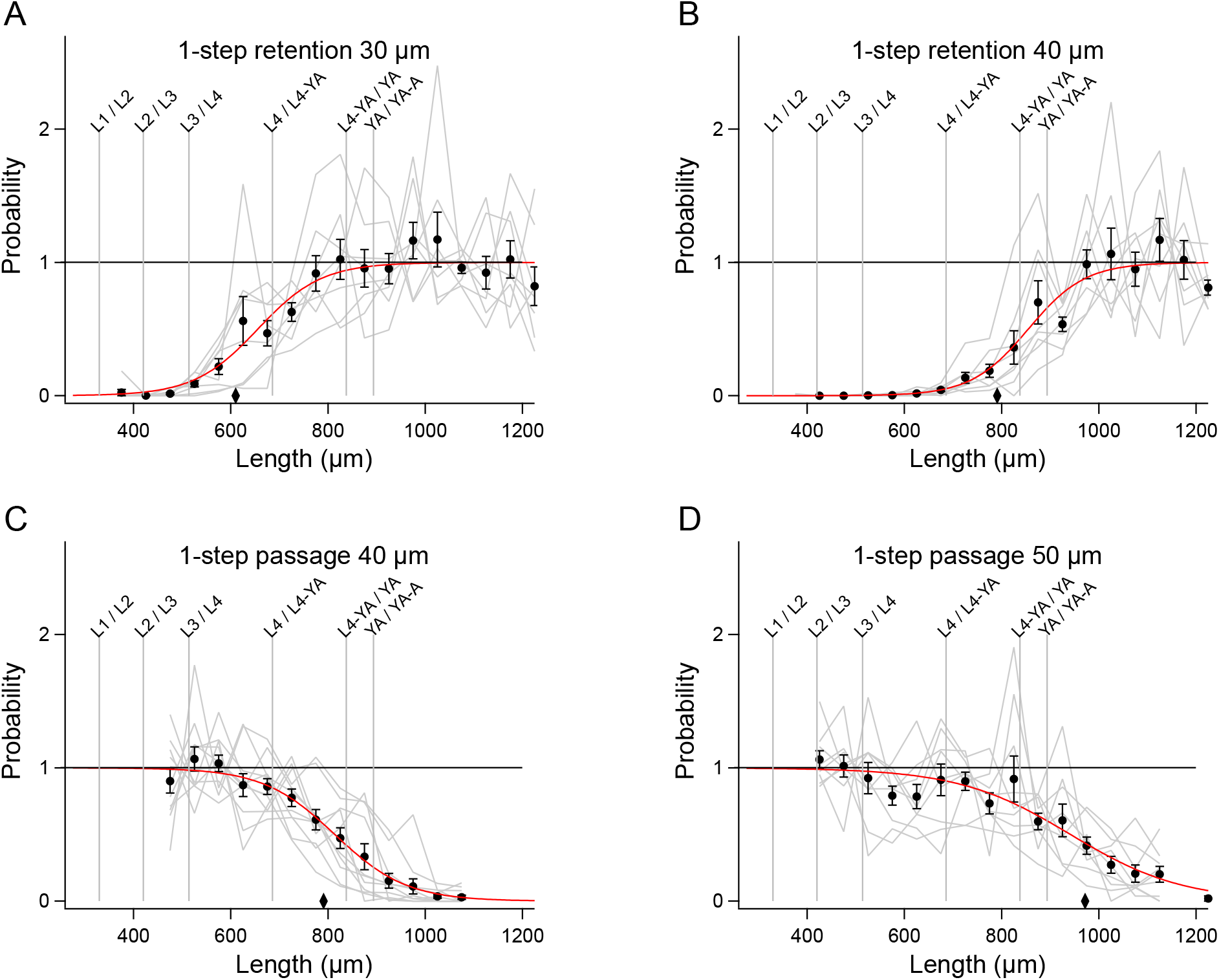
1-step filter functions extracted from the data in Fig 3. A-D. *Gray traces* show the value of the ratio on the right-hand side of equation 2 for each run, after multiplication by the constant *C* (see text), such that the mean of data points in the asymptotic region is 1. Number of data points in the asymptotic region used to compute *C* is given in S1 Table. *Black symbols* indicate means of gray traces at each length. Error bars, SEM. *Red traces* are fits of equation 3 or 4 to the means in each panel. Diamonds indicate the length of worms having a diameter equal to the mesh size.

To validate the filter functions, we retroactively predicted the length distributions of filtered worms by multiplying the mean unfiltered distribution 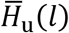 by the filter function. The predicted filtered distributions closely matched the actual filtered distributions, 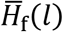 (Fig 5A-D). We conclude that the filter functions accurately represent the filtration process despite considerable run-to-run variability in the data (Fig 4, gray traces).

**Fig 5.**
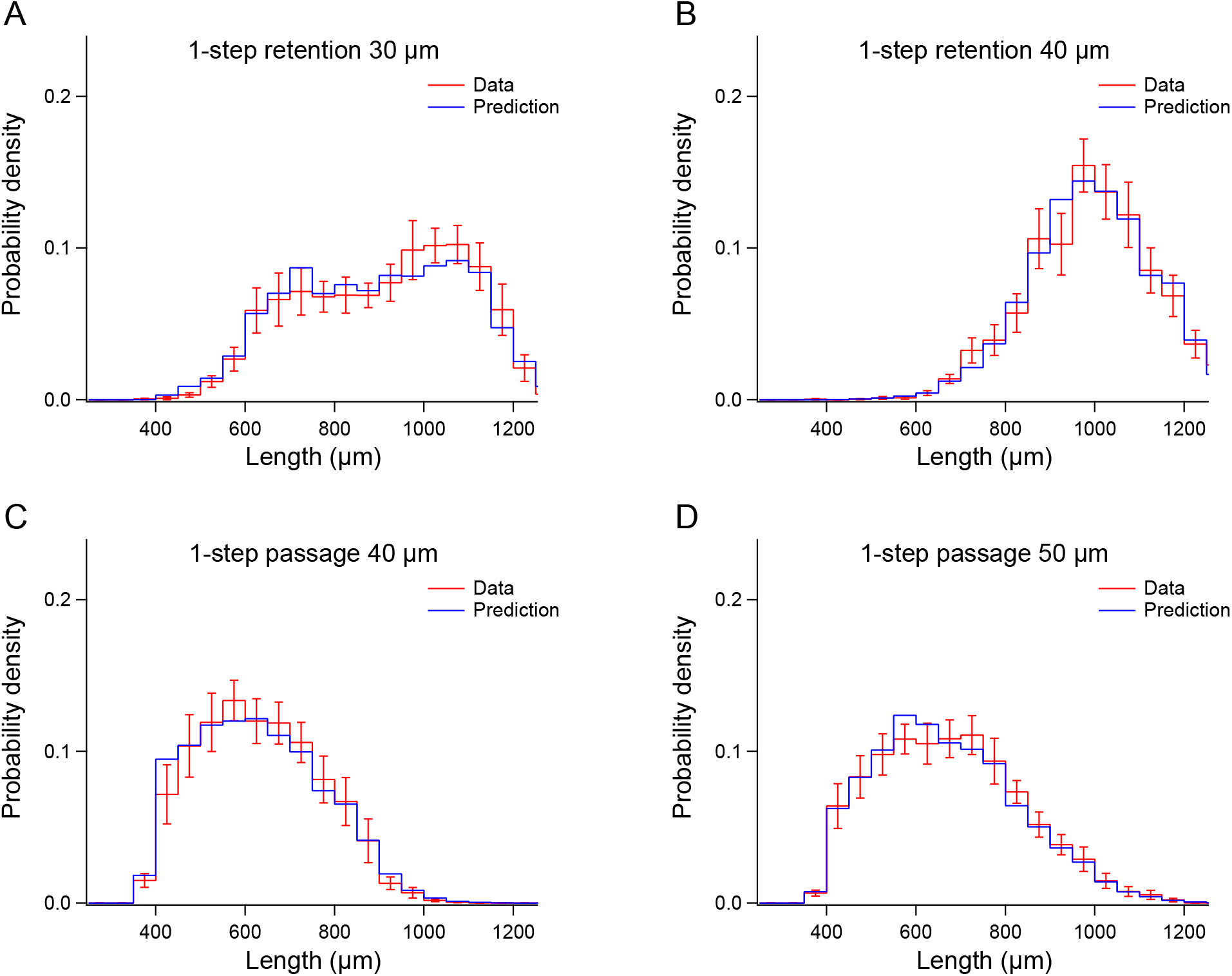
Validation of 1-step filter functions. A-D. In each panel, the unfiltered length distributions 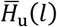 in the corresponding panel of Fig 3 were multiplied by the filter functions in the corresponding panel of Fig 4, yielding the *blue trace* (Prediction). The *red trace* (Data) shows the actual filtered distribution 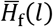 from the corresponding panel of Fig 3. Error bars, SEM.

### Comparison with sieving theory

We next tested the hypothesis that filtration of C. *elegans* conforms to the predictions of ideal sieving theory in two distinct experiments. The first experiment involved a multi-step filtration procedure in which a filtered fraction is re-filtered one or more times through the same filter. In the simple example of a two-step retention procedure, the retained fraction is filtered a second time. By basic laws of probability, the probability that a particle is retained after the first *and* the second filtration step is the product of the retention probabilities for each step, *p*(*x*)^2^. In the case of *n* filtration steps, the retention probability is *p*(*x*)^*n*^.

The second experiment considered the relationship between the probabilities of retention and passage. In an ideal sieving process, particles are either passed or retained by the sieve, and all retained particles are recovered from it. Mathematically, *p*_r_(*x*) + *P*_p_(*x*) = 1, where *p*_r_(*x*) is the probability of retention, *p*_p_(*x*) is the probability of passage. In the case of the 40 μm cell strainer, for which we obtained filter functions for both filtration modes, we found that these functions did not add to unity (Fig 6, dashed trace). In particular, there was a significant probability of a worm being neither passed nor retained, which is computed as 1 – [*p*_r_(*l*) + *P*_p_(*l*)] (Fig 7, blue trace). This is a second respect in which filtration of worms by cell strainers does not conform to sieving theory.

**Fig 6.**
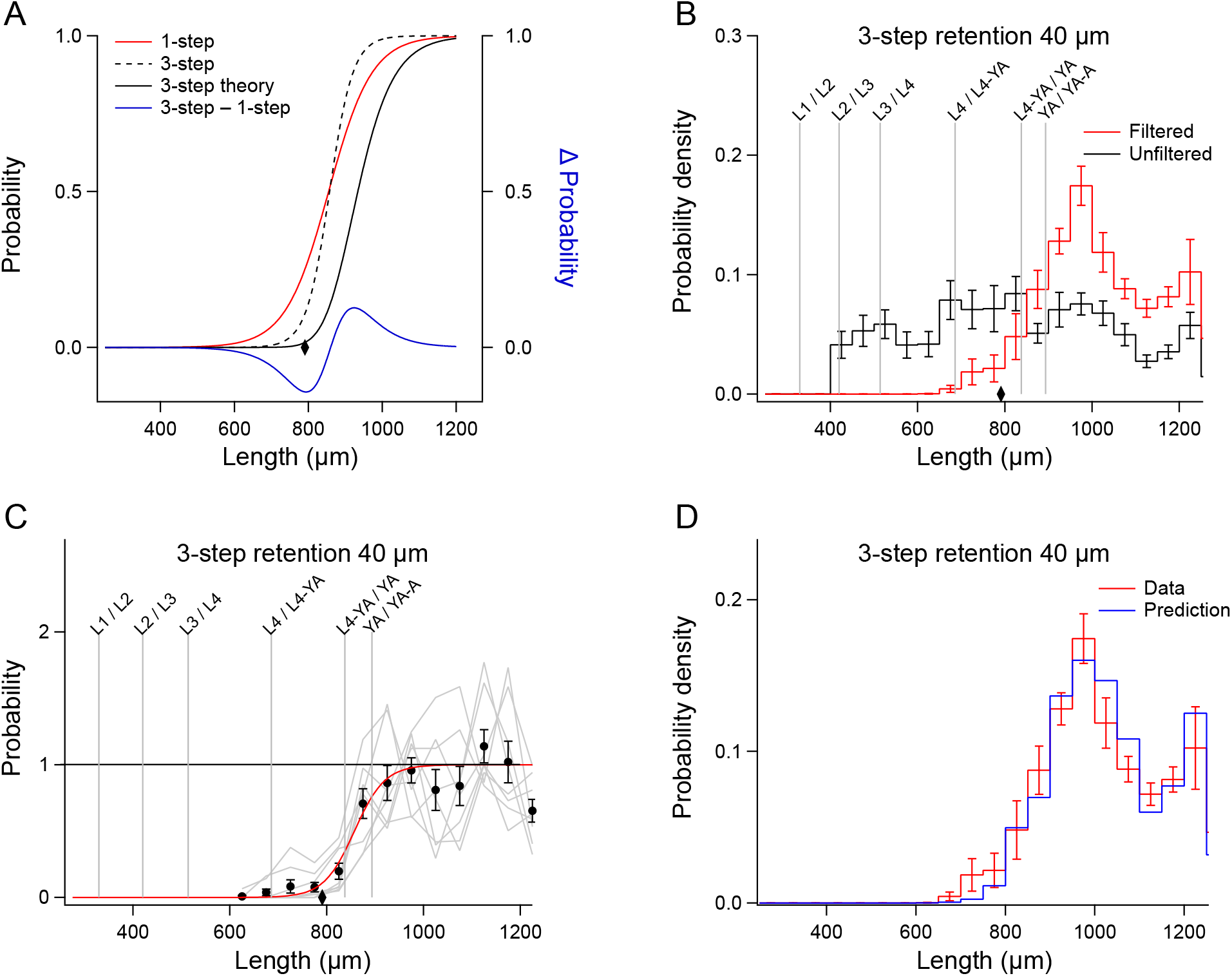
Non-ideal properties of 3-step filtration. A. Comparison of theoretical and actual filter functions. The *red trace* is 1-step retention, 40 μm filter function from Fig 4B. The *black trace* is the 1-step filter function raised to the third power. The *dashed black trace* is the actual three-step filter function obtained by the analysis shown in C. The *blue trace* is the difference between the 1-step and 3-step filter functions. B. Filtered distributions, 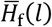 (*red trace*), and unfiltered distributions, 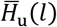 (*black trace*). The diamond indicates the length of worms having a diameter equal to the mesh size. Error bars, SEM. C. Filter function for 3-step filtration. *Gray traces* show the value of the ratio on the right-hand side of equation 2 for each run, after multiplication by the constant *C* (see text) so that the mean of data points in the asymptotic region is 1. *Black symbols* indicate means of gray traces at each length. Error bars, SEM. The *red* trace is the fit of equation 3 to the mean data. Number of extreme data points in the asymptotic region used to compute *C* is given in S1 Table. D. Validation of the 3-step filter function. The unfiltered length distribution 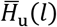 in B was multiplied by the filter function in C, yielding the *blue trace* (Prediction). The *red trace* (Data) shows the actual filtered distribution 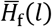 in B.

**Fig 7.**
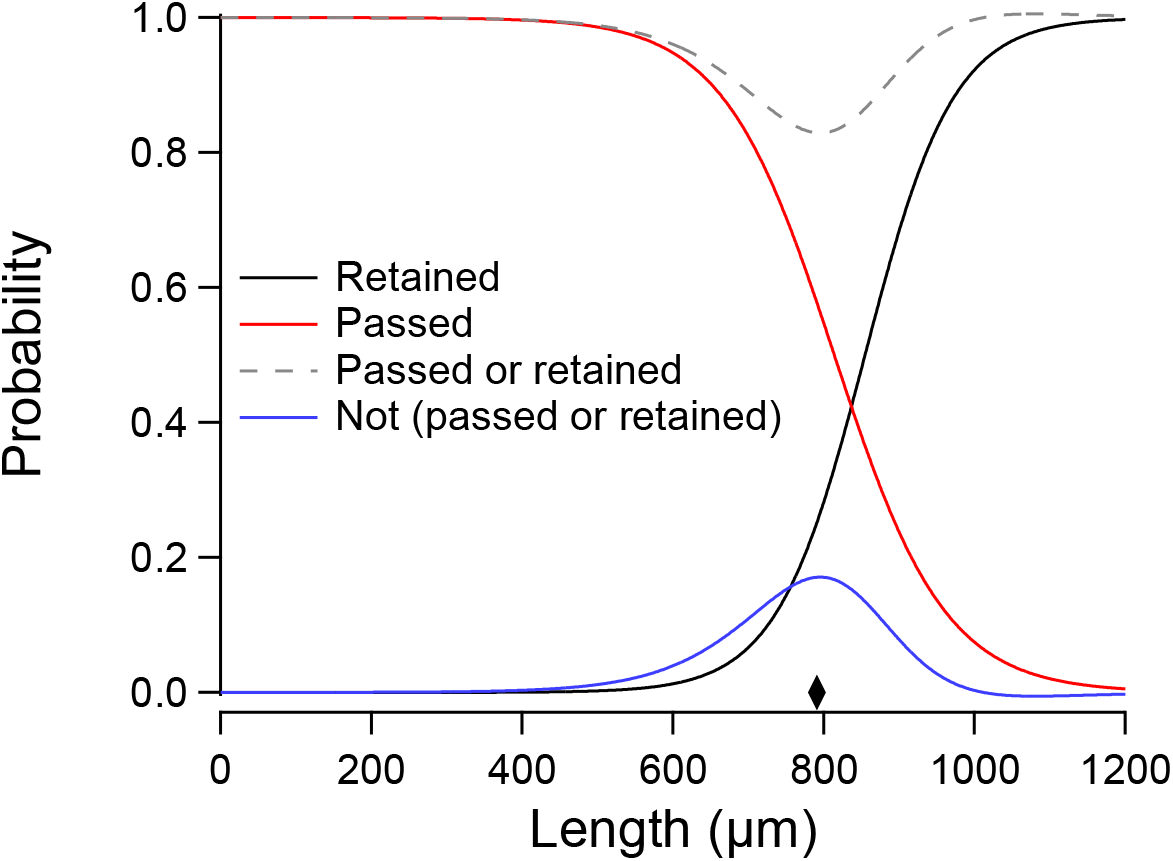
Non-ideal properties of 1-step filtration. The *black* and *red traces* are the filter functions for 1-step retention, 40 μm (Fig 4B) and 1-step passage, 40 μm (Fig 4C), respectively. In the ideal case, the sum of filter functions for retention and passage by the same filter should equal 1. However, for the 40 μm cell strainer it does not (*gray dashed trace*). The probability of being neither passed nor retained (*blue trace*) is maximal at 795 μm, a length corresponding to a width of 40.2 μm (*diamond*) which is close to the mesh size. This finding suggests that worms having a width approximately equal to the mesh size have a higher probability of being captured by the mesh than wider or narrower worms.

However, it is notable that the probability of being neither passed nor retained reached its maximum at a length corresponding to a width almost equal to the mesh size (40 μm, Fig 7, diamond). This correspondence predicted that some worms are be captured by the mesh. To test this prediction, we filtered a starting population using the 1-step retention, 40 μm method. We then back flushed the strainer in the usual way (Materials and Methods), submerged the strainer in M9 buffer, and inspected the mesh under a stereomicroscope. We found a significant number of worms whose heads were caught in the mesh, with their tails free to engage in normal swimming movements (S1 Video). Furthermore, the fact that some worms are caught by the strainer means that the fraction of worms recovered in retention mode actually represents worms that were retained *and* subsequently released by the filter.

The finding that worms can be caught by the strainer helps explain the fact that sieving theory does not correctly predict the outcome of multi-step filtration (Fig. 6A). As noted above, with successive filtration steps, the probability of retaining and releasing longer worms increases whereas the probability of passing shorter worms decreases. A possible explanation for increased retention of longer worms is that the number of mesh openings of the right size to capture such worms is reduced as these openings progressive become more occupied by captured worms. A possible explanation for decreased retention of shorter worms is that, being narrower, they are captured less tightly during initial filtration steps, and are eventually ejected from the mesh by fluid pressure in subsequent filtrations.

These findings highlight the utility of theoretical filter functions in two key respects. Without knowledge of the 40 μm strainer’s retention function, it would have been impossible to characterize the non-ideal properties of this strainer. On the other hand, we have shown that filter functions can be predictive. In particular, the 40 μm retention and passage functions correctly predicted trapping, another finding that would not have been possible by simple inspection of filtered and unfiltered length histograms. Thus, the filter functions provided a means of elucidating the mechanics of worm filtration using cell strainers. This knowledge will be useful when designing novel filtration protocols, including filtration of other species of nematodes.

### Filtration as an alternative to bleach synchronization

Guided by our knowledge mechanics of worm filtration by cell strainers, we sought to devise a filtration protocol that might serve as an alternative to synchronization. As proof of concept, we sought to obtain populations of worms in the range dominated by developmental stages L4 and YA. In the N2 reference strain, worms in this range vary in width from 35 to 42 μm [7]. Cell strainers closest to this range are 30 μm and 40 μm strainers. We tested three filtration methods, each based on a 40 μm strainer to exclude older adult worms and a 30 μm strainer to capture L4s and YAs. *L4–YA Method 1* (Fig 2F) was a 2-step approach: a 40 μm strainer in passage mode, followed by re-filtration of the recovered worms with a 30 μm strainer in retention mode (Fig 8A). *L4–YA Method 2* (Fig 2G) was a 1-step approach using a 40 μm strainer stacked on a 30 μm strainer (40/30); worms were recovered from the 30 μm filter (Fig 8B). *L4–YA Method 3* (Fig 2H) was a 2-step approach using the 40/30 stack, in which worms recovered from the 30 μm strainer in step 1 were filtered a second time (Fig 8C). Length histograms of filtered worms were well fit by gaussian functions, parameters of which are given in Table 2. L4–YA Methods 2 and 3 yielded distributions with peaks in the target range, whereas the peak of L4–YA Method 1 was shifted toward longer worms (Fig 8 and Table 2). Differences in the dynamics of fluid flow between stacked and non-stacked strainers might account for this shift.

**Fig 8.**
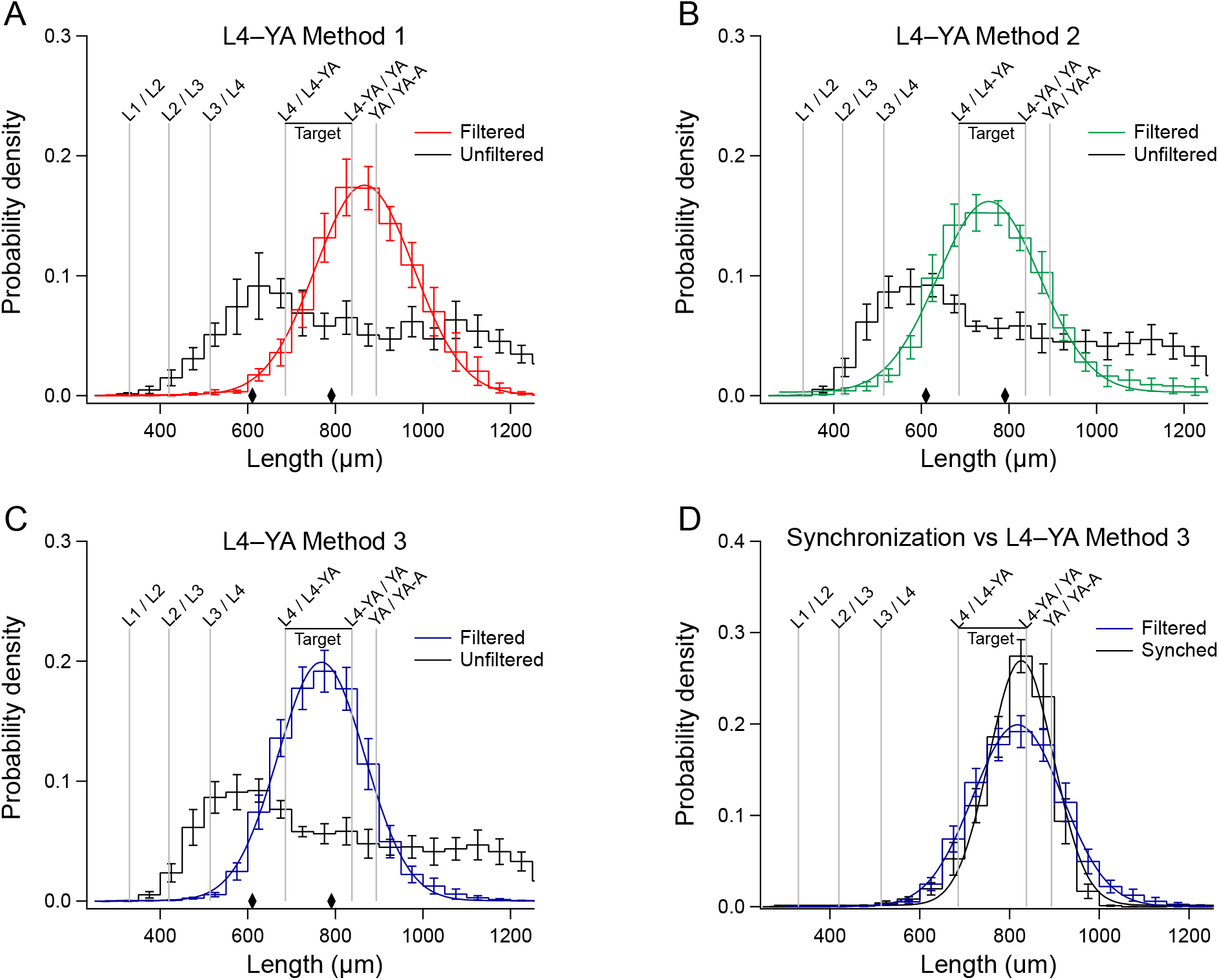
Enrichment by filtration for particular developmental stages. The goal was to enrich for developmental stages L4 and YA (*Target*) utilizing a 40 μm strainer and a 30 μm cell strainer in three distinct protocols (L4–YA Methods 1-3, Fig 2). A-C. Filtered distributions, 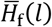 (*colored traces*), and unfiltered distributions, 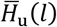 (*black traces*). *Diamonds* indicate the length of worms having a diameter equal to the two mesh sizes. A. L4–YA Method 1, filtration by a 40 μm strainer in passage mode, followed by re-filtration of the recovered worms with a 30 μm strainer in retention mode. B. L4–YA Method 2, 1-step filtration by a 40 μm strainer stacked on a 30 μm strainer. C. L4–YA Method 3, 2-step filtration in which worms recovered from the 30 μm filter in Method 2 were passed through the stack a second time. D. Comparison of synchronization (*black trace*) and L4–YA Method 3 (*blue trace*). To facilitate comparison of synchronization and L4–YA Method 3, the distribution of the latter was shifted to the right by 50 μm. A-D. Error bars, SEM.

Methods of enrichment by size-sorting can be compared in terms of throughput, yield, and purity. The literature on *C. elegans* microfluidic size-sorting devices defines throughput as the number of worms that can be sorted per minute (wpm). Throughput using the Biosort is approximately 180 wpm (Erik Andersen, personal communication). In the case of microfluidic devices, throughput when sorting from mixed-stage populations varies from 4 wpm [9, 13] to 200 wpm [10], depending on device design. In this study, the highest throughput was obtained using L4–YA Methods 2 and 3, for which we used 10 culture plates per run. To compute throughput, we estimated the number of worms in 10 plates (see Materials and Methods), divided this number by the processing time for each run, and computed the within-method mean across runs. Throughput ranged from approximately 7,000 to 14,000 wpm, exceeding the maximum throughput of previous size sorting methods (200 wpm) by a factor of 35 to 70.

Mean yields for L4–YA Methods 2 and 3 are given in Table 2. These values are lower-bound estimates because it was impractical to measure every worm in the filtered population. However, these estimates suggest that both methods can generate hundreds to thousands of worms per run.

Purity was defined as the proportion of worms whose lengths were within the target zone (Fig 8A-D), which extends from 686 μm (L4 / L4-YA) to 838 μm (L4-YA / YA). Purity on each run was computed as the integral within the target zone of probability densities *H*_u_(*l*) and *H*_f_(*l*). Purity for synchronized populations was computed in a similar manner, with one exception. The peak of the mean probability density function for synchronized worms fell within the target zone but was not centered on it (Fig 8D). Therefore, before computing purity values for each synchronization each run, we centered its histogram by shifting it 39 μm to the left. This approach eliminated the spurious reduction in purity that would have resulted from the slight mismatch between the synchronized distribution and the target zone.

Mean purity for unfiltered populations associated with individual runs of L4–YA Methods 2 and 3 was 0.191 (± 0.080 std. dev., *n* = 15 replicates). Mean purities for filtered and synchronized populations are given in Table 2. Mean purities for L4–YA Methods 2 and 3, whose filtered distributions were well aligned with the target zone (Fig 8B,C), exceeded those of unfiltered populations (L4–YA Method 2 vs. Unfiltered: *t* = 7.00, df = 11.89, *p* = 1.50 × 10^-5^; L4–YA Method 3 vs. Unfiltered: *t* = 9.46, df = 11.51, *p* = 9.01 × 10^-7^); this comparison shows that these methods had a significant effect on purity. Furthermore, the purity of L4–YA Method 3 was greater than that of L4–YA Method 2 (*t* = 2.26, df = 11.98, *p* = 0.043), indicating that the second step of filtration improved performance substantially. We next obtained the length distribution for bleach synchronized worms (Fig 8D). The purity of synchronization (Materials and Methods) was greater than that of L4–YA Method 3 (*t* = 2.50, df = 13.28, *p* = 0.026). The peak of the distribution of synchronized worms was in the target range, as expected (Table 2), but not centered on it. To facilitate visual comparison of synchronization and L4–YA Method 3, we aligned the peaks of the two distributions by shifting the distribution of L4–YA Method 3 to the right by 50 μm. Inspection of this graph emphasizes the fact that although the purity of synchronization is greater than the purity of L4–YA Method 3, the latter may be sufficient, in practical terms, especially in applications where accessibility, high throughput, and yield are priorities.

## Discussion

We combined theory and experiment to investigate the use of cell strainers to enrich for particular developmental stages of *C. elegans* by filtration. We found that this method of filtration can be described by a filter function that worm length to the probability of recovery (retention or passage), with length being a proxy for developmental stage. Recovery in both modes was well-described by logistic functions, which are rotationally symmetric about the point at which probability is 0.5. This symmetry suggests that passage and retention may be governed by a single process, possibly how tightly worms of a given length, hence a given width, fit within the mesh. Cell strainers are a generic laboratory commodity, with identical meshes regardless of manufacturer. Thus, our main findings are not manufacturer dependent.

We found that filtration of worms by cell strainers is consistent with ideal sieving theory [25] in some, but not all, respects. The correspondence we observed between actual and retroactively predicted distributions of filtered worms is consistent with theory (Fig 5A-D, 7D). This finding is significant because it means users can predict the outcome of 1-step filtration based on their particular starting distributions, which is highly variable across strains, experimenters, and laboratories. There were two main inconsistencies. First, in the ideal case, the filter function for multistep filtration through the same filter is equal to the 1-step filter function raised to the power of the number of filtration steps. This was not the case for 3-step retention, 40 um filtration (Fig 6A). Such inconsistencies are not surprising. Second, in an ideal sieving process, particles are either passed or retained by the sieve; none are trapped by it. This was not the case for 1-step retention, 40 μm filtration, as there was a significant probability that worms were not released by the filter (Fig 6). Indeed, inspection of the filter after washing showed that some worms remained trapped in the mesh (Supplemental Video 1). Ideal sieving theory is based on rigid, spheroid particles whereas worms are flexible, elongated, and capable of self-movement. In practical terms, these inconsistencies mean that the outcome of a multistep filtration procedure cannot be predicted *a priori;* it must be determined experimentally.

*C. elegans* enrichment methods can be compared along multiple dimensions, including throughput, purity, accessibility, yield, and detrimental physiological effects. Fig 9 compares currently available methods along the first three dimensions. Throughput data fall into two domains (low and high) whereas the purity data fall into three domains (low, intermediate, and high). Accessibility has two levels, low (grey) and high (blue). Microfluidic chips and Biosort occupy the low-throughput, high-purity, low-accessibility sector. These methods are best suited to applications in which high purity is sufficiently important to justify the effort and expense required to achieve it. Filtration by L4–YA Methods 2 and 3 occupies the high-throughput, intermediate-purity, and high-accessibility sector. These methods are best suited to applications that require large numbers of worms that can be obtained quickly and are compatible with intermediately levels of purity. Synchronization occupies the low-throughput, intermediate-purity, high accessibility sector. This method is best suited to low throughput applications in which concerns about the detrimental effects of bleach are outweighed by the modest increase in purity over L4–YA Method 3.

**Fig 9.**
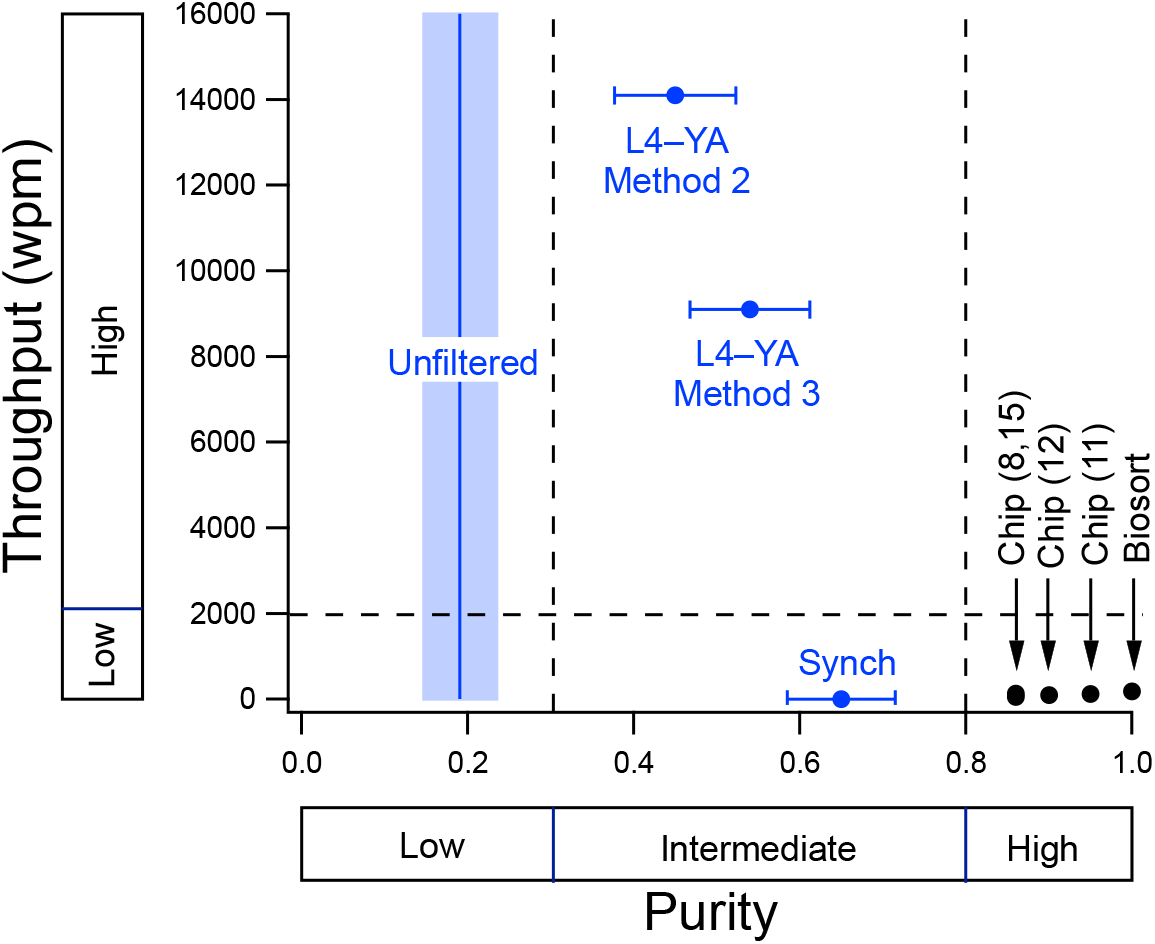
Comparison of enrichment methods along the dimensions of throughput, purity, and accessibility. Throughput is defined as the number of worms processed per minute. Purity is defined as the proportion of worms whose lengths are within the target zone (L4 and YA) when sorting from a mixed-stage population. The mean purity of unfiltered worms is indicated by the blue vertical line. Accessibility (high, *blue symbols*; low, *black symbols*) is defined as the inverse of the magnitude of resources, facilities, and training required to make use of the method. Microfluidic chips and Biosort occupy the low-throughput, high-purity, low-accessibility sector. The L4–YA Methods 2 and 3 and synchronization occupy the high-throughput, intermediate-purity, and high-accessibility sector. Error bars and shading, 95% CI.

### Limitations

Several limitations should be kept in mind when designing new filtration protocols using cell strainers. (1) Filter functions obtained here are limited to filtering strains that closely resemble N2 in width and length at each developmental stage. These functions will likely not apply to strains carrying mutations that alter worm diameter (e.g., “dumpy” (*dpy*) and “multi-vulva” (*muv*) mutants) or length (e.g., “small” (*sma*) and “long” (*lon*) mutants). Nor will they necessarily apply to populations that have undergone treatments, such as drugs or altered temperature, that change worm size.

These functions also might not apply to mutants (e.g., “uncoordinated” (*unc*) mutants) or treated populations having reduced swimming ability, as this defect could alter the orientation in which worms contact and subsequently interact with the mesh (Supplemental Video 1, blue arrow at *t* = 4 sec). It will therefore be necessary to experiment with a range of different mesh sizes for sorting such strains. (2) Cell strainers are generally available in mesh-size increments of 10 μm, which can limit the precision of the sorting process, especially in the case of stacked filtration to enrich for a particular larval stage. However, mesh fabrics from which custom strainers could be constructed are available in finer increments (S2 Table), opening the possibility of replacing the mesh of a cell strainer with a mesh of intermediate mesh size. (3) When enriching for particular larval stages, the purity of filtered populations in this study was less than the purity of synchronized populations (Table 2). This problem could be mitigated by increasing the number of filtration steps, as L4–YA Method 3 (2 steps) had significantly higher purity than L4–YA Method 2 (1 step).

### Modifications

Several modifications of the present protocols could expand the utility of the filtration approach. Stacked filtration based on other pairs of mesh sizes could be utilized to enrich for stages other than L4-YA. For example, a 50/40 stack could be used to enrich specifically for YA worms used in DNA injections. Microfluidic sorting devices that enrich for multiple developmental stages simultaneously have been demonstrated [7, 11, 12, 15]. This functionality could be replicated with cell strainers by increasing the number of strainers in a stack. For example, a 40/30/25/20 stack could be used to enrich simultaneously for larval stages L4 and L3 in addition to the L4-YA mixture obtained from the 40/30 stack (Fig 8). Another modification is to increase filtration throughput and yield by custom fabrication of larger strainers, which could be as simple as replacing the bottom of plastic drinking cups with mesh fabric.

### Applications

The simplest filtration method uses a single cell strainer. This approach could be utilized to enrich for adult worms from mixed-stage populations by using a strainer that passes all stages except adults. The stringency of this type of enrichment could be improved by multi-step filtration over the strainer, as shown in Fig 7A. This approach could facilitate behavioral, genetic, and genomic experiments by enabling the use of very large populations of adults. Single-strainer filtration could also be used either to select or exclude morphological mutants when mixed with wild type worms. For example, *dpy* or *muv* mutants could be selected or excluded using a cell strainer that passes wild type worms but not these mutants. As the *dpy* locus is frequently used as a balancer for maintaining strains carrying lethal mutations, this approach could increase sample sizes in studies of lethal genes, one of the largest gene classes in *C. elegans*. We have also shown that stacked-strainer filtration can be used to enrich for particular larval stages and that the purity of this approach can be increased by repeated filtration. This approach could facilitate developmental studies that focus on a wide range of stage-specific genetic, physiological, and behavioral mechanisms.

## Acknowledgements

Maggie Kerner contributed technical assistance to the research.

## Supporting information

**S1 Fig.**
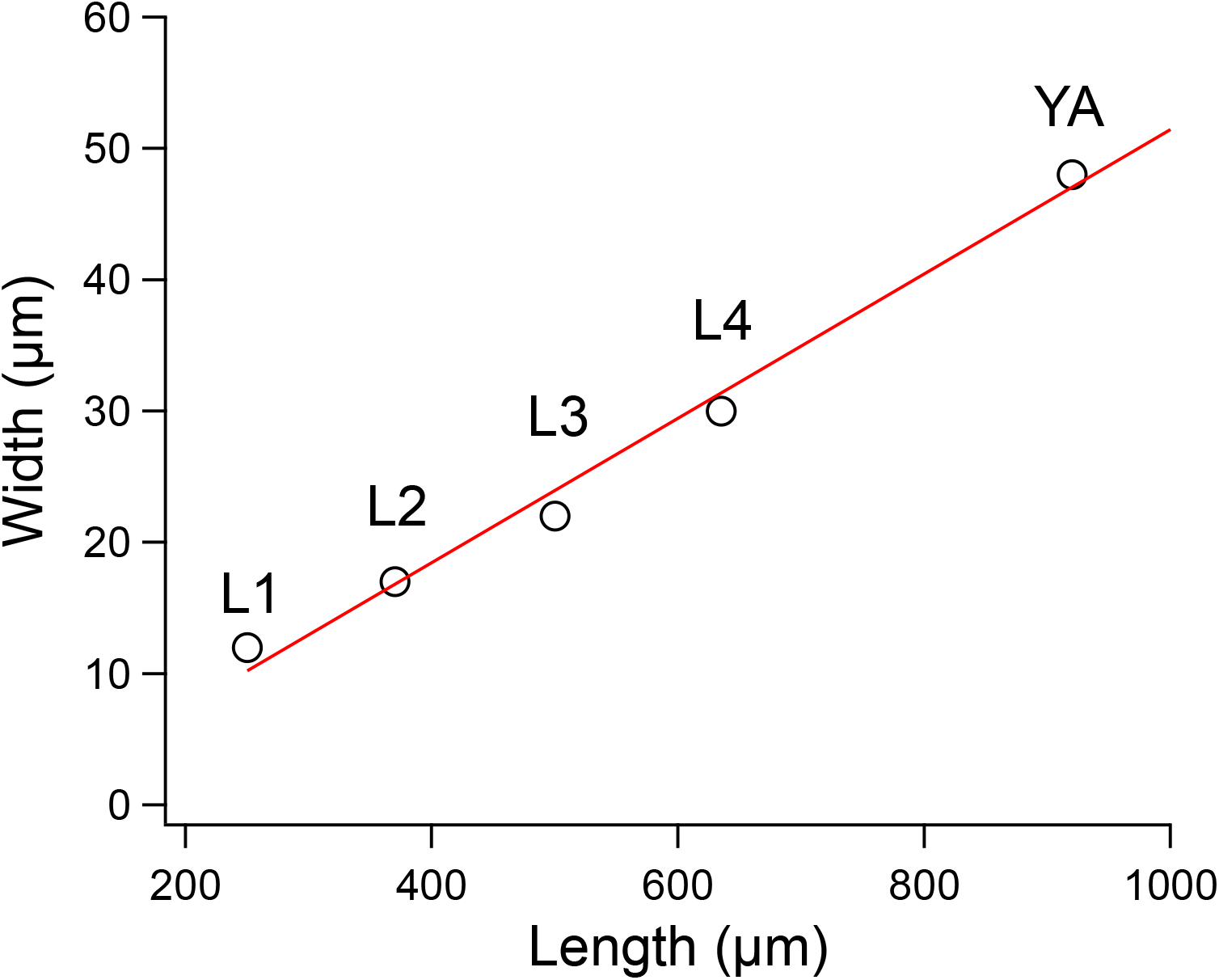
Width-length curve. *Widths* of each stage were taken from measurements in Table 1 of Atakan et al. [7]. *Lengths* of each stage were the center of the length ranges that define each stage according to data in WormAtlas [23]. The data are fit by the equation *w* = 0.055*l* — 3.53, where *w* is width and *l* is length.

**S2 Fig.**
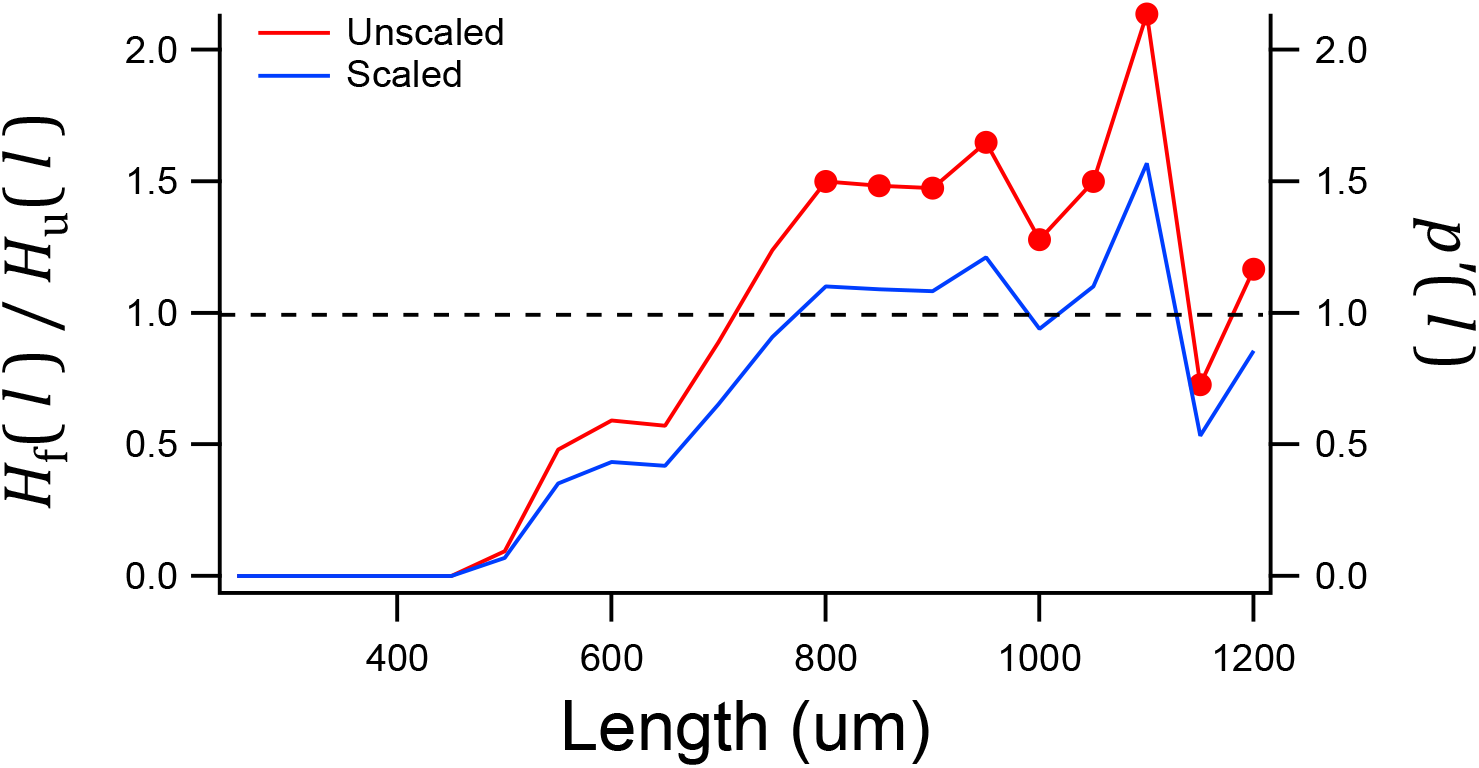
Scaling procedure used to constrain limiting values of *p′*(*l*). *Red trace*, the ratio *H*_f_(*l*)/*H*_u_(*l*) is plotted against length for one run of 30 μm retention. *Blue trace*, the same data after scaling *y*-values by 1/*X_n_*, with n = 9. The value of n was chosen by trial and error to optimize the fit of the logistic function to the mean of all runs in the 30 μm retention data set.

**S1 Table 1.**
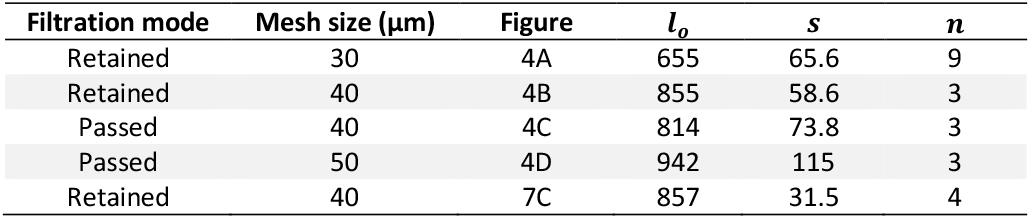
Filter-function parameters for fits of equations 3 and 4 to the data in the indicated figures. Column *n* contains the number of values used to define the asymptotic region of the data during the fitting procedure.

**S2 Table 2.** Examples of available nylon mesh sizes. ^a^Component Supply, Sparta, TN, 38583 USA. ^b^pluriSelect, El Cajon, CA, 92020 USA. ^c^Funakoshi, Tokyo, Japan.

| Mesh fabric<br>($\mu\text{m}$)<sup>a</sup> | Cell strainers<br>($\mu\text{m}$)<sup>b</sup> | Cell strainers<br>($\mu\text{m}$)<sup>c</sup> |
| --- | --- | --- |
| 7 | 1 | 25 |
| 10 | 5 | 40 |
| 15 | 10 | 70 |
| 18 | 15 |  |
| 25 | 20 |  |
| 31 | 30 |  |
| 38 | 40 |  |
| 40 | 50 |  |
| 44 | 60 |  |
| 52 | 70 |  |
| 56 | 85 |  |
| 60 | 100 |  |
| 62 |  |  |
| 64 |  |  |
| 70 |  |  |
| 80 |  |  |
| 85 |  |  |
| 105 |  |  |

**S1 Video. Worms caught by the cell-strainer mesh.** After 1-step retention, 40 μm filtration, the strainer was flushed in the usual way to recover worms from the top surface of the mesh. It was then submerged in M9 buffer. Inspection of the mesh on a stereomicroscope revealed worms whose heads were caught in the mesh. Their tails exhibited normal swimming movements (*red arrows*). The video also shows a swimming worm that gets caught in the mesh (*t* = 15 sec, *blue arrow*), suggesting that worms actively interact with the mesh in some cases.

## Derivation of the logistic function

Let *p*(*x*) be the probability of a favorable outcome in a binary, categorical process, where *x* is a continuous variable that predicts *p*. By definition, the odds of a favorable outcome are the ratio of the probability of a favorable outcome to the probability of a non-favorable outcome

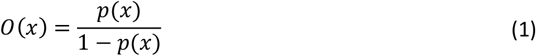

In logistic regression, the parameters of a linear equation in *x*, such as *y* = (*x* – *μ*)/*s*, are adjusted to fit the probability data *p*(*x*) after transformation as log of the odds of a favorable outcome.

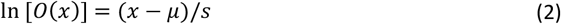

which is equivalent to

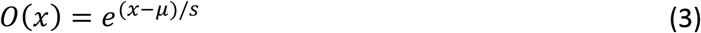

or

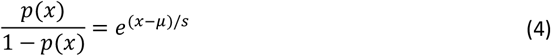

Solving for *p*(*x*) gives gives the familiar form of the logistic function

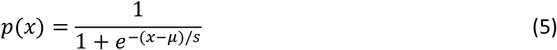

